# CNNM1 is Involved in Spermatogonial Stem Cell Maintenance in the Mouse

**DOI:** 10.1101/2025.10.20.683346

**Authors:** Irfan khan, Pradeep G Kumar

## Abstract

Male reproductive development starts early during embryogenesis with the formation of spermatogonial stem cells (SSCs) from their neonatal precursors, the gonocytes. Previously, we reported high levels of CNNM1 expression in the testicular germ cells of neonatal mice, which were maintained in the SSCs of pubertal and adult mice. CNNM1 exhibited a progressive decline in expression levels as the SSCs progressed through the spermatogenic differentiation sequence. However, the specific functional role of CNNM1 during spermatogenesis remains elusive. Here, we studied the expression pattern of Cnnm1 during gonocyte to spermatogonia transition (GST), and expression levels were found to be higher at the initial establishment phase of SSCs marked with a higher rate of cell proliferation. GDNF-mediated self-renewal/proliferation of the C18-4 spermatogonial cell line enhanced the expression of CNNM1. Overexpression of CNNM1 in C18-4 spermatogonial cell lines resulted in the upregulation of genes involved in cell proliferation, nucleic acid metabolism, male germ cell development and various pathways involved in cell cycle regulation, whereas CNNM1 knockdown altered the cell cycle progression. Based on the expression analysis and proteome profiling, we conclude that CNNM1 regulates and maintains the self-renewal, proliferation, and survival of mouse spermatogonial cells.

**Graphical Abstract:** 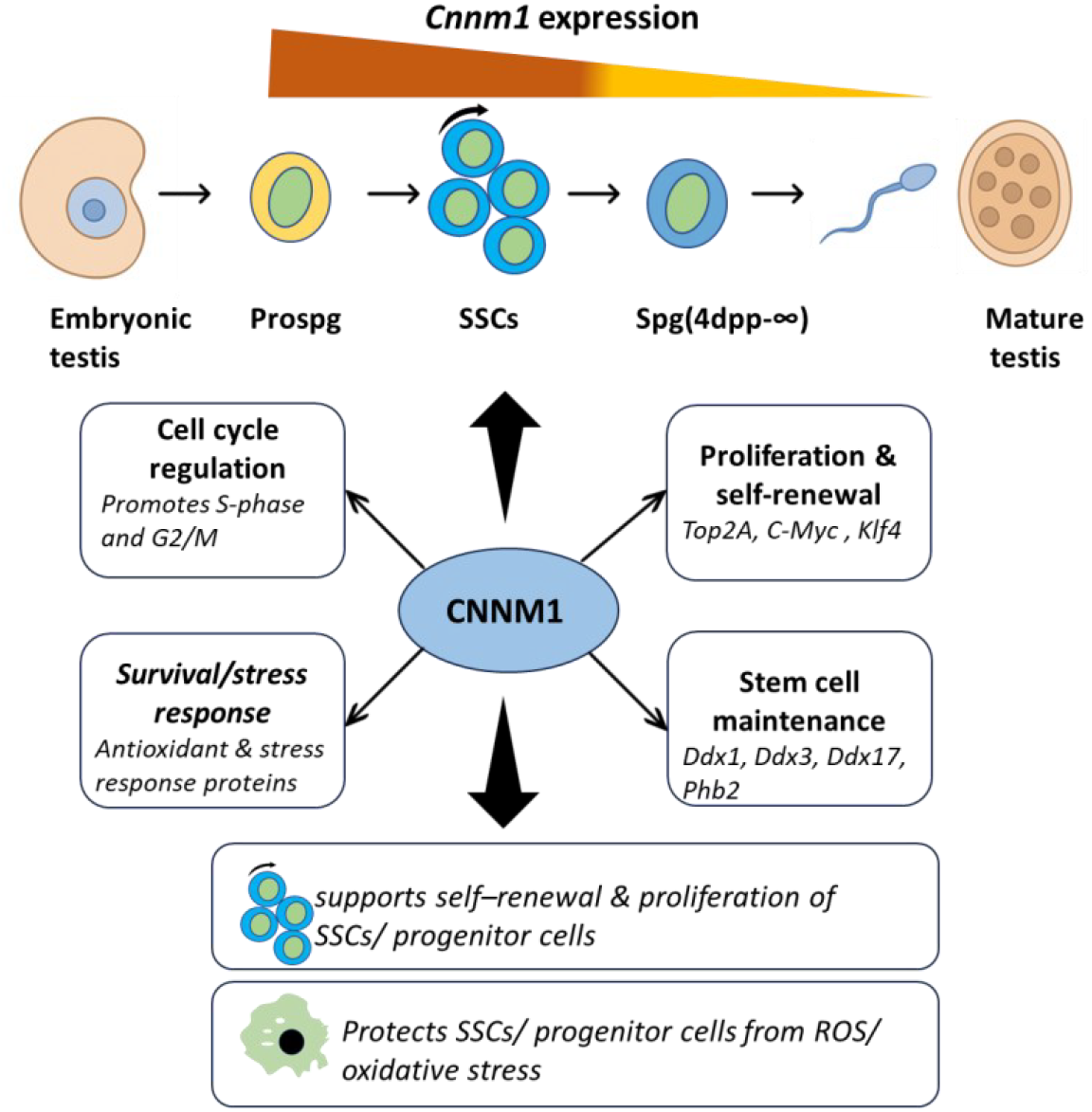

## Introduction

Mammalian spermatogenesis is a classic adult stem cell–dependent process that requires systematised differentiation and timely coordinated cellular changes ^1, 2^. Spermatogonial stem cells (SSCs) are a group of adult stem cells in the testis, capable of self-renewal to maintain their stem cell pool and differentiation to produce mature spermatozoa ^1^. Thus, SSCs are the foundation of continuous spermatogenesis and male fertility. In the testis, prospermatogonia are derived from primordial germ cells (PGCs) after the sex specification event as precursors of SSCs. In mice, the first appearance of a foundational population of SSCs occurs at approximately 3-4 dpp ^1^. The quintessential undifferentiated SSC population can simultaneously divide asymmetrically to produce differentiated spermatogonia and to maintain a reservoir of undifferentiated SSCs through self-renewal throughout spermatogenesis ^3^. Despite their importance, the low number of SSCs in the testis has greatly hindered our ability to understand the biology of SSCs ^4^. Due to the complex microenvironment contributed by the surrounding testicular somatic cells, it has been difficult to understand the molecular mechanisms and signalling pathways that regulate the SSC fate determination towards self-renewal or differentiation ^1, 4^. Identifying the molecular modulators that determine the SSC fate is vital in the development of diagnostic and therapeutic interventions for male infertility.

*Cnnm1* is a putative cell cycle regulator highly expressed in the brain and the testis ^5^. Previous studies had shown that CNNM1 regulates cell cycle, proliferation, survival, and tumor development in prostate cancer and hepatocellular carcinoma ^5-7^. Data from our laboratory had identified CNNM1 as an aberrantly expressed protein in male factor infertility (unpublished). In mice, CNNM1 expression was higher during the neonatal stages than in adults. Interestingly, testicular expression levels of the CNNM1 dropped after postnatal day 8. Immunohistochemical analysis of mice testis showed perfect co-localization of CNNM1 and SSC markers, and differentiation of SSC led to the loss of its expression ^5^. In this study, we investigated the potential role of CNNM1 during the early stages of male germ cell specification and gonocyte to spermatogonia transition in mice. *Cnnm1* was abundantly expressed during the proliferative phase of SSC establishment. Overexpression of CNNM1 in C18-4 spermatogonial cell line resulted in proteome-wide changes that could affect metabolism, cell proliferation and stress responses in male germ cells.

## Results

### Expression of *Cnnm1* during mouse gonocyte to spermatogonia transition

Previous studies had shown high expression of CNNM1 in neonatal mouse testes and its strong localization in SSCs in adult mice testis, implying its possible role in establishing the SSC population. To address this aspect, we evaluated the expression profile of *Cnnm1* throughout the gonocyte to spermatogonia transition during fate reprogramming and establishment phases of SSC development. Gonads from E18.5 embryos and neonatal testis from 0 dpp to 8 dpp animals were analysed. Prospermatogonia present in E16.5 and E18.5 testes enter mitosis during the establishment phase with a marked increase in proliferation and self-renewal of these cells between 2 dpp and 3 dpp to build the foundational SSC pool. A quantitative reverse transcription polymerase chain reaction (qRT-PCR) analysis demonstrated a significant increase in the levels of *Cnnm1* in 3 dpp testis **(Fig. 1A)**, followed by a decline at 4 dpp, suggests the potential role of *Cnnm1* in self-renewal and proliferation of the SSC population. These results implied that CNNM1 expression might be essential after the sex specification of PGCs.

**Figure 1.**
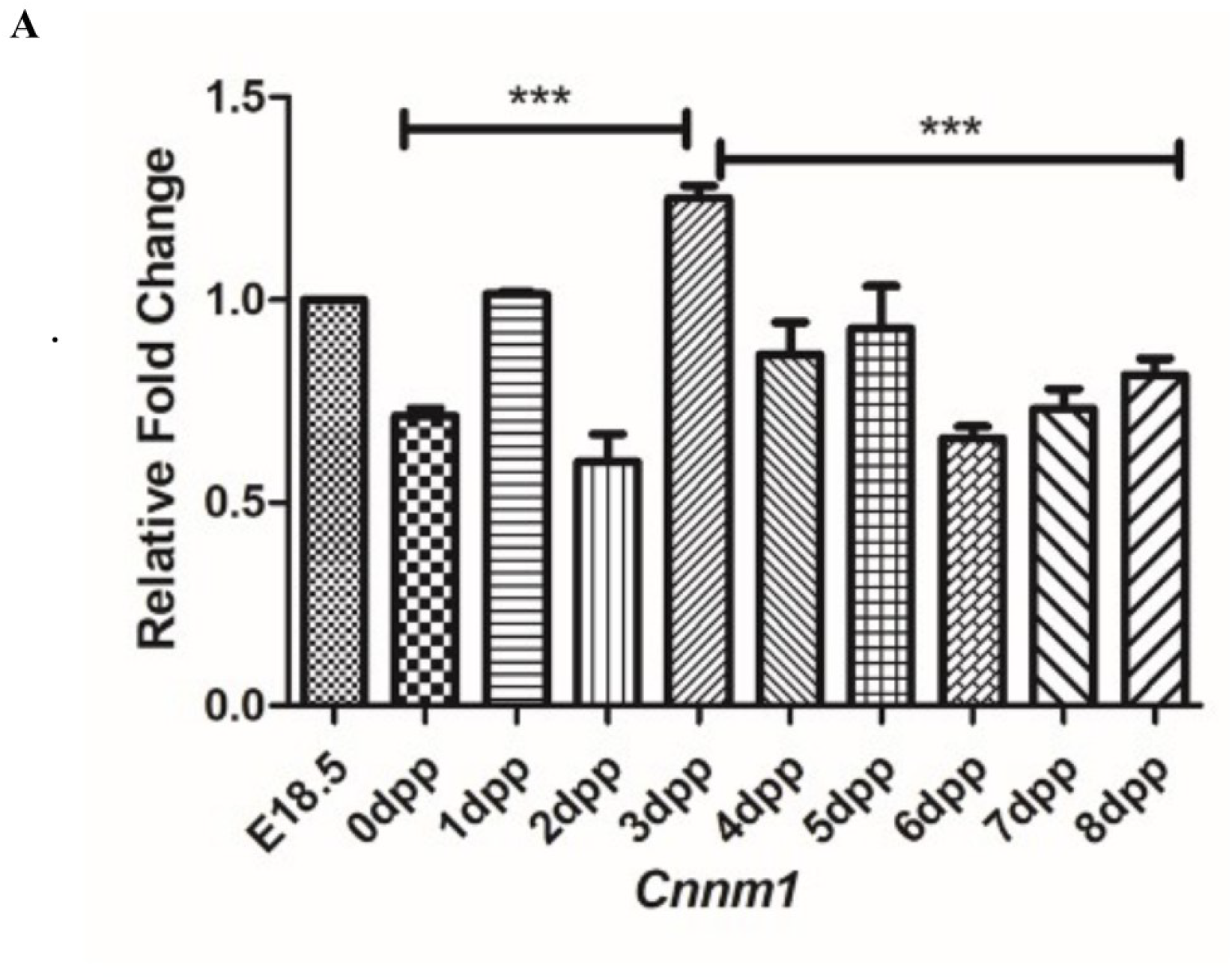
Expression of *Cnnm1* during gonocyte-to-spermatogonia transition in mice. Quantification of *Cnnm1* transcript levels across developmental stages (A). Bars represent mean ± s.e.m. Statistical analysis was performed using GraphPad Prism; P < 0.05 was considered significant. *Gapdh* served as control E = embryonic, dpp = days postpartum.

### Glial cell-derived neurotropic factor (GDNF) - mediated self-renewal of spermatogonial cells upregulated the CNNM1 expression

GDNF, ligand for GFRA-1, is a self-renewal factor that plays a critical role in determining the fate of SSCs. To test whether *Cnnm1* is associated with the proliferation and self-renewal of mouse SSCs, C18-4 spermatogonial cells were stimulated with GDNF (100ng/ml). This resulted in a significant increase in *Cnnm1* expression both at protein **(Fig. 2A)** and transcript levels **(Fig. 2B)**. Immunofluorescence results confirmed the upregulation CNNM1 expression in C18-4 cells treated with GDNF compared to untreated control cells **(Fig. 2C)**.

**Figure 2.**
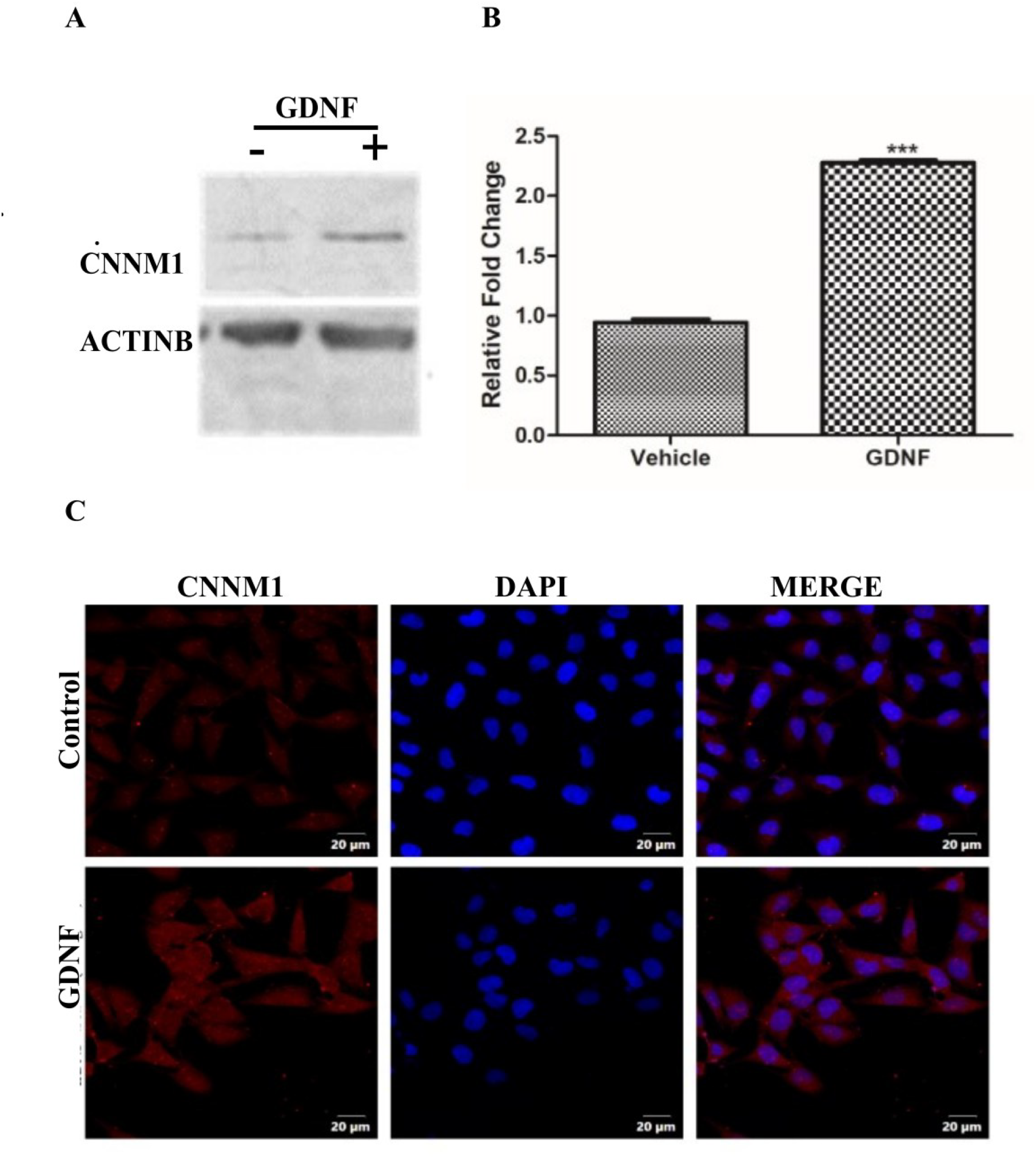
GDNF treatment upregulates CNNM1 expression in C18-4 cells. Western blot analysis of CNNM1 expression in GC1 and C18-4 cells treated with GDNF (100ng/ml) (A), Bars represent the expression level of *Cnnm1* transcript in GDNF-treated C18-4 cells. (B), Immunofluorescence study of GDNF-treated C18-4 cells, CNNM1 expression (a, d). DAPI (b, e) and Merge (c, f); Bar = 20 μm (C). Statistical analysis was performed using GraphPad Prism software, and p<0.05 was calculated.

### Generation of the *Cnnm1* overexpression construct

*Cnnm1* complementary DNA (cDNA) amplified from neonatal mouse testis was cloned into the pEGFP N1 vector **(Fig. S1, B)** and was used for transient transfection of C18-4 cells. Immunofluorescence analysis of *Cnnm1*-pEGFPN1 and pEGFPN1 transfected cells probed with CNNM1 antibody showed a heavy colocalization within the nuclei of cells overexpressing CNNM1**(Fig S1, A)**.

### CNNM1 overexpression led to an increase in cell cycle progression and proliferative capacity of spermatogonial cells

The C18-4 cells transfected with pEGFPN1 and *Cnnm1*-pEGFPN1 were subjected to Fluorescence-activated cell sorting (FACS), 24 h after transfection. GFP-positive cells were sorted by FACS, and cell cycle analysis was performed by Hoechst 33342 staining **(Fig. 3, A and B)**. It was observed that there was a reduction in G1-phase population and an increase in G2/M phase populations of *Cnnm1*-pEGFPN1 cells compared to pEGFPN1 control cells **(Fig. 3, C)**. Furthermore, RT-PCR analysis showed an upregulation of Top2A expression with CNNM1 overexpression **(Fig. S1, C)**.

**Figure 3.**
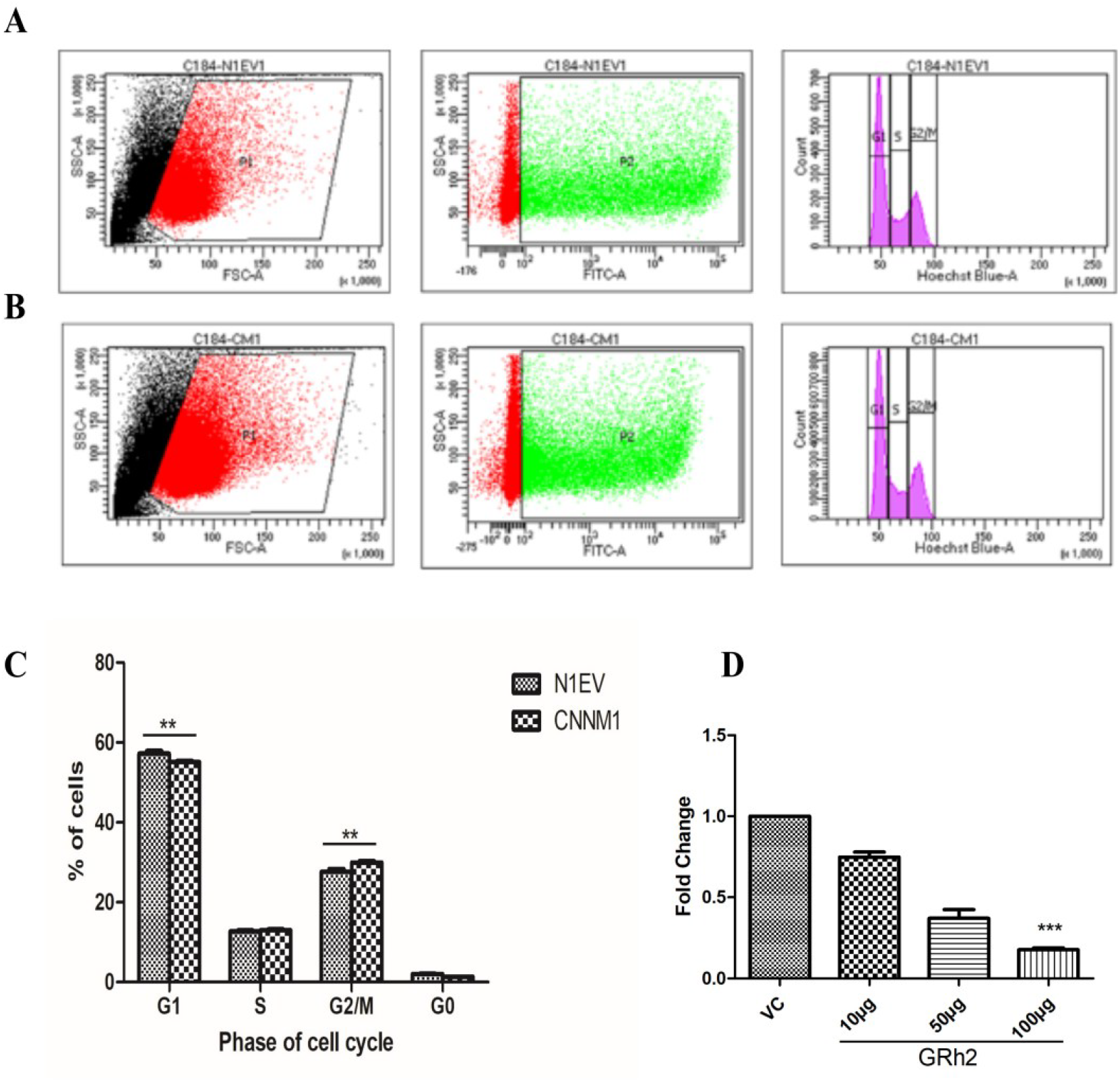
CNNM1 overexpression promotes cell cycle progression in spermatogonial cells and is downregulated by a pro-apoptotic agent. Cell cycle analysis of GFP-positive C18-4 cells 24 h after transfection with pEGFPN1 (A) or CNNM1-pEGFPN1 (B), stained with Hoechst 33342 and corresponding bar graphs show increased S-phase and G2/M-phase populations in CNNM1-overexpressing cells compared with control (B). Dose-dependent downregulation of *Cnnm1* upon treatment with Ginsenoside Rh2 (G-Rh2, 10, 50, 100 µg/ml). demonstrate a significant decrease relative to vehicle-treated controls (D). Statistical analysis was performed using GraphPad Prism software, and p<0.05 was calculated.

### Proapoptotic agent downregulates *Cnnm1* expression in spermatogonial cells

Since *Cnnm1* overexpression altered the cell cycle progression and upregulated the proliferative marker Top2A **(Fig. S1, C)**, we evaluated the influence of already reported proapoptotic factor Ginsenoside Rh2 on *Cnnm1* expression in C18-4 cells. The cells were treated with three gradient concentrations (10, 50 and 100 µg) of G-Rh2 (Ginsenoside Rh2) for 48h. G-Rh2 treatment brought about a There was a significant dose-dependent decrease in the expression of *Cnnm1* when compared with the vehicle control **(Fig. 3, D)**.

### Global proteome analysis reveals association of cell cycle and metabolic processes with CNNM1 overexpression in spermatogonial cells

A quantitative proteomic analysis was performed using liquid chromatography in tandem with mass spectrometry to identify differentially expressed proteins between CNNM1-overexpressed and control cells. A total of 989 proteins were identified, of which 215 proteins were upregulated (fold change >1.5), while 13 proteins were downregulated (fold change 0.5). A list of differentially expressed proteins from the proteome profile of CNNM1 overexpressed cells, along with fold change, is provided **(Table S1)**.

The biological processes of differentially expressed proteins in control and CNNM1 overexpressed C18-4 cells were identified using gene ontology (GO) annotation and enrichment analyses. In gene ontology, 253 biological processes were assigned to the 170 genes that showed significant changes. Based on the number of attributed genes, the top biological processes of altered proteins after CNNM1 overexpression are cellular process (GO:0009987, 102 hits), metabolic process (GO:0008152, 62 hits), biological regulation (GO:0065007, 37 hits), and localization (GO:0051179, 22 hits), and developmental process (GO:0032502, 5 hits), **(Fig. 4, A)**. Furthermore, secondary and tertiary level annotation revealed significantly altered cellular processes, including upregulation of cell differentiation, cell cycle, reproduction and cell morphogenesis processes in CNNM1 overexpressed C18-4 cells.

**Figure 4.**
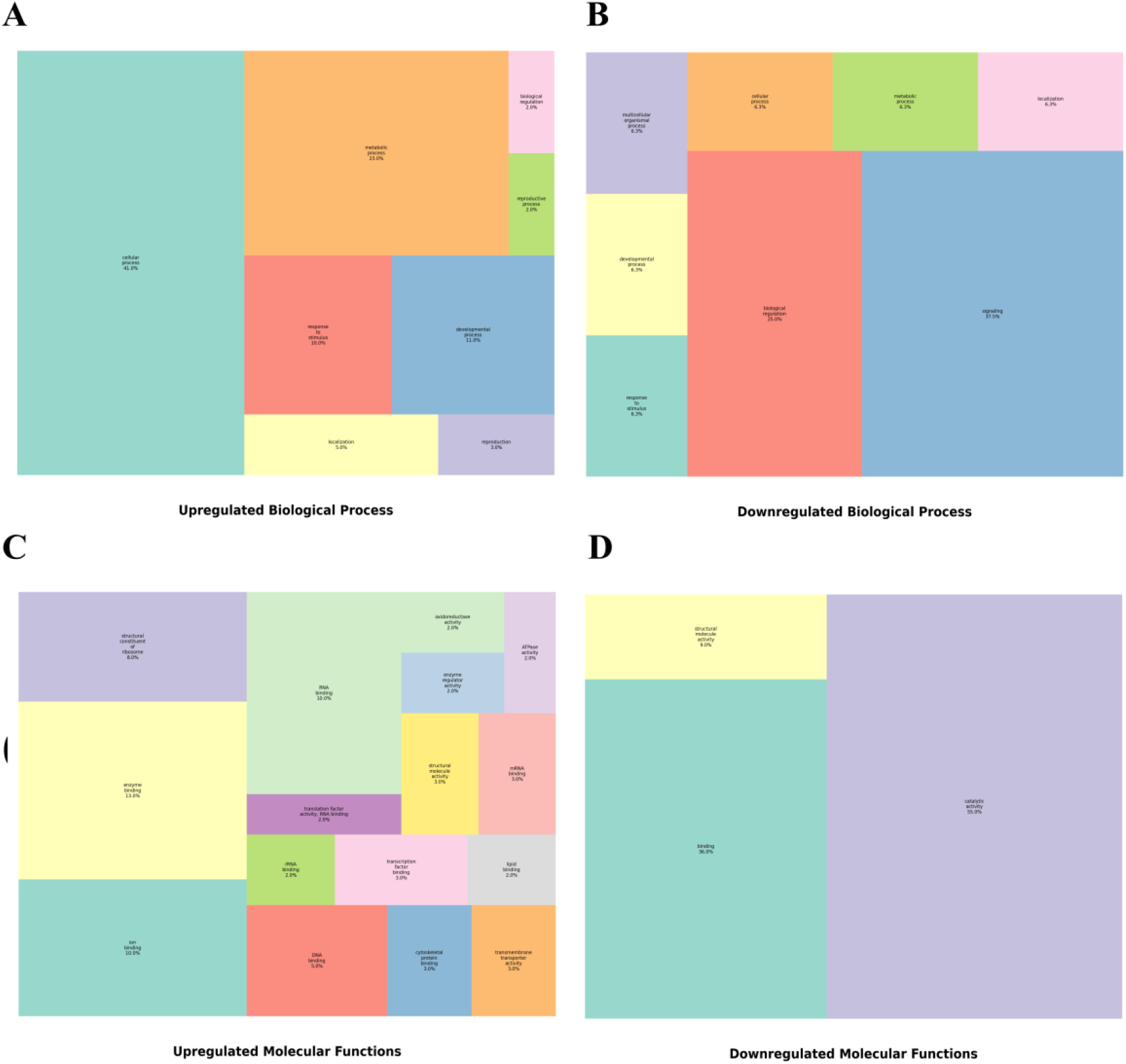
Proteomic profiling of CNNM1-overexpressing spermatogonial cells. Gene ontology (GO) enrichment of upregulated proteins in CNNM1-overexpressing cells; top biological processes include cell cycle, proliferation, reproductive and developmental processes (A). Downregulated processes include cellular metabolism and nitrogen compound processing (B). GO analysis of molecular functions of upregulated (C) and downregulated (D) proteins, highlighting catalytic and binding activities. Proteins with fold change > 1.5 are considered to be upregulated and fold change < 0.5 are considered to be downregulated.

Utilizing the same criteria, enrichment analysis of downregulated proteins suggested 16 biological processes that were downregulated in the CNNM1 overexpressed C18-4 cells. Among the 16 biological processes, cellular process (GO:0009987,6 hits), metabolic process (GO:0008152, 4 hits) are the top processes to be downregulated **(Fig. 4, B)**. Secondary and tertiary level annotation revealed cellular processes such as cellular metabolic process, cellular nitrogen compound metabolic process, and phosphorus metabolic process to be downregulated.

### Global proteome analysis reveals association of ion binding and ligase activity functions with CNNM1 overexpression in spermatogonial cells

GO and enrichment analysis were carried out to identify the molecular functions of altered proteins. GO annotation showed that the 213 genes have generated a total of 219 molecular function hits of which, binding proteins (GO:0005488, 94 hits), catalytic activity (GO:0003824, 52 hits), structural molecule activity (GO:0005198, 32 hits) and transporter activity (GO:0005215, 11 hits) were the top functional hits of altered proteins **(Fig. 4, C)**. Since protein binding function was one of the top hits, further secondary and tertiary level annotation revealed ion binding, RNA binding and cytoskeletal binding proteins were significantly upregulated in CNNM1 overexpressed cells.

On the other hand, the top functional hits of downregulated proteins were catalytic activity (GO:0003824, 6 hits) and binding (GO:0005488, 4 hits) **(Fig. 4, D)**. Since protein binding function was one of the top hits, further secondary and tertiary level annotation revealed cyclo-ligase activity, ligase activity, forming carbon-nitrogen bonds, amide binding, ligase activity and nucleotide binding were downregulated in CNNM1 overexpressed cells.

### Global proteome analysis reveals association of Wnt signalling pathway with CNNM1 overexpression in spermatogonial cells

GO annotation carried out by Panther analysis to identify the key pathways associated with upregulated proteins in CNNM1-overexpressed C18-4 cells generated 128 pathway hits. Wnt signalling pathway, ribosome biogenesis and PI3 Kinase signalling pathway were the major pathways which were upregulated. The de novo purine biosynthesis pathway (P02738) was found to be associated with downregulated proteins **(Fig. S2)**.

### Network analysis of altered proteins reveals association of male gamete development processes with CNNM1 overexpression in spermatogonial cells

GOnet network analysis was performed for the altered proteins specifically enriched in the reproductive process in CNNM1 overexpressed cells. Several biological processes associated with male germ cell development, including the mitotic cycle, germ cell development and proliferation, reproductive developmental process, and spermatogenesis to be upregulated upon overexpression of CNNM1 **(Fig. 5, A)**.

**Figure 5.**
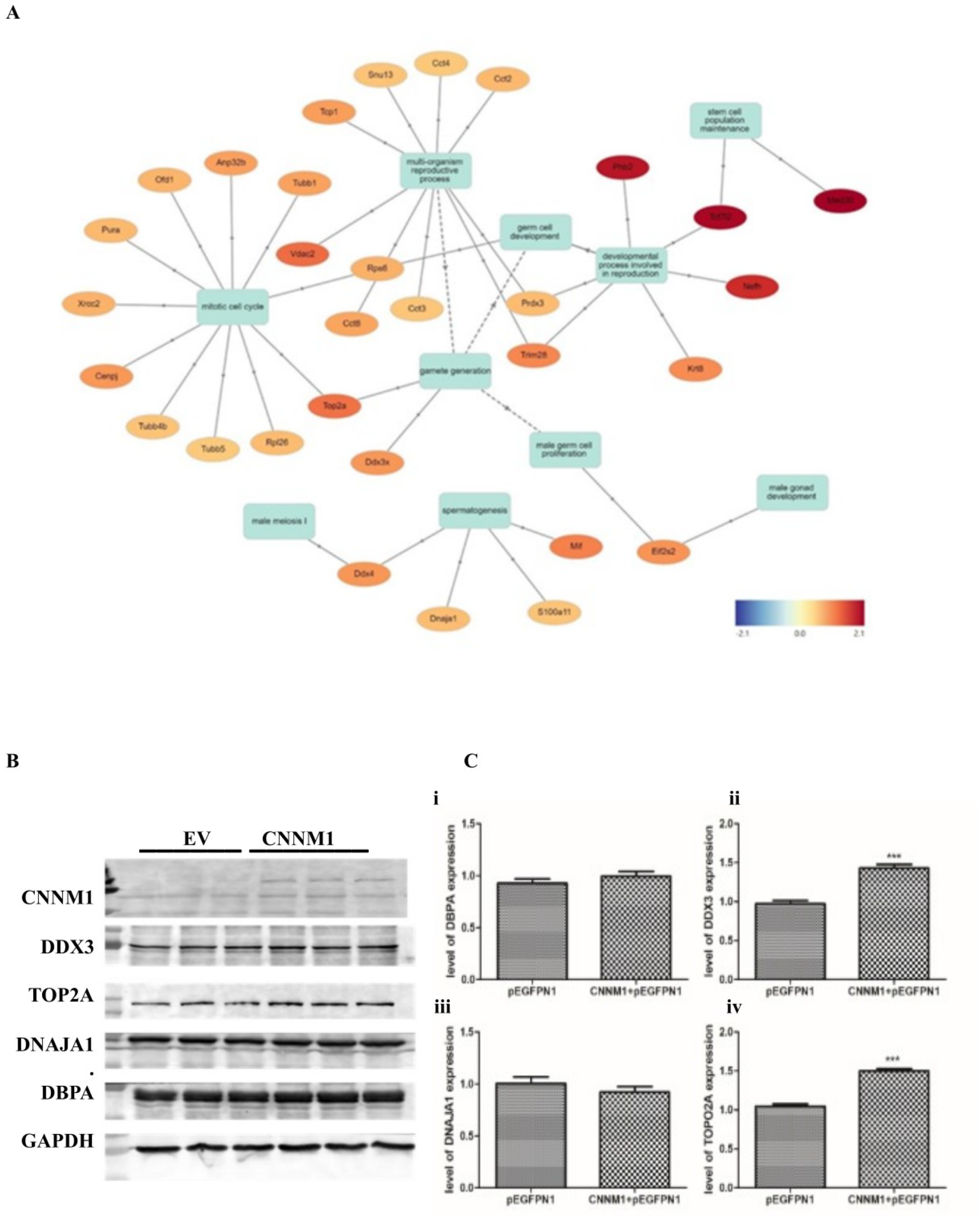
CNNM1 overexpression modulates proteins associated with spermatogonial proliferation and male germ cell development. GONet network analysis of proteins upregulated upon CNNM1 overexpression, showing enrichment in biological processes related to the mitotic cell cycle, germ cell proliferation, and spermatogenesis (A). Representative immunoblots showing expression of DDX3, TOP2A, DNAJA1, and DBPA in C18-4 spermatogonial cells transfected with control pEGFP-N1 (EV) or CNNM1-pEGFP-N1 constructs (CNNM1 OE); GAPDH was used as a loading control (B). Densitometric quantification of protein bands from three independent experiments. A significant increase in DDX3 and TOP2A expression was observed, whereas DBPA and DNAJA1 levels remained unchanged (C, i-iv). Statistical analysis was performed using GraphPad Prism software, and p<0.05 was calculated.

### CNNM1 overexpression upregulated the proteins involved in cell proliferation and stem cell maintenance proteins in spermatogonial cells

Overexpression of CNNM1 in spermatogonial cells resulted in the upregulation of proteins associated with cell processes and the cell cycle. Therefore, from our protein panel, we focused on the expressions of genes involved in these processes, which showed more than > 1.5 upregulation. We validated the proteins DNA Topoisomerase II Alpha (TOP2A, 170 kDa), DEAD-box helicase 3 (DDX3, 73kDa), DNA-binding protein A (DBPA, 40kDa) and DnaJ homolog subfamily A 1 (DNAJ1, 45 kDa) by immunoblot analysis and compared the expressions of these proteins with control pEGFPN1-transfected cells **(Fig. 5B)**. Western blot analysis confirmed significantly elevated expression of DDX3 and TOP2A in CNNM1-overexpressing cells, which supports our proteomic data **(Fig. 5C i-iv)**. DDX3 is an essential gene involved in stem cell proliferation and maintenance by regulating apoptosis and the cell cycle ^8^. It is also crucial for the proliferation of mouse spermatogonia with inferred evidence in azoospermic men ^9^. TOP2A is an essential gene involved in proliferation. These results suggest that CNNM1 might play an essential role in spermatogonial cell proliferation and maintenance.

## Discussion

Previous studies have reported that CNNM1 is highly expressed in the neonatal testis, restricted to the SSC population, and downregulated upon differentiation ^5^. Therefore, in the present study, we aimed to shed light on the functional role of CNNM1 in SSC establishment and maintenance.

We analysed Temporal expression profiling revealed that *Cnnm1* transcript levels remained largely unchanged during PGC specification and PGCLC induction from ES cells using a protocol adapted from Hayashi et al.^10^, suggesting that Cnnm1 is not essential during the earliest stages of germline commitment. However, expression increased significantly from E13.5, coinciding with sex determination and the initiation of testicular differentiation. This elevation persisted throughout the various phases of GST (E13.5–8 dpp), encompassing fetal mitotic-prospermatogonia or M-gonocytes (E13.5), transitional 1 prospermatogonia or T1 gonocyte (E16.5-0 dpp) and transitional 2 prospermatogonia or T2 gonocyte (0 dpp to 3 dpp) and in the neonatal stage (4 dpp to 8 dpp) ^11, 12^. Notably, *Cnnm1* expression peaked during the highly proliferative phase associated with SSC establishment (E18.5–3 dpp) and declined thereafter (4–8 dpp), concurrent with SSC maturation and the emergence of differentiating A-type spermatogonia. These data suggest that CNNM1 is functionally linked to the proliferative expansion of the foundational SSC population rather than later differentiation.

To further explore this function, we employed the C18-4 mouse spermatogonial cell line, which retains key molecular and phenotypic characteristics of SSCs ^13^. Treatment of C18-4 cells with GDNF—a key paracrine regulator of SSC self-renewal resulted in a marked induction of Cnnm1 expression. GDNF acts through the GFRα1/c-RET receptor complex to activate downstream PI3K/AKT and ERK/MAPK pathways, which are essential for SSC survival and proliferation ^14^. The GDNF-mediated upregulation of Cnnm1 therefore implicates CNNM1 as a downstream effector or target of self-renewal signaling in spermatogonial cells. Previous reports showed that GDNF enhances proliferation in C18-4 cells via Ras/ERK1/2–FOS signaling ^15^ further support this interpretation.

Recent studies have implicated CNNM1 in regulating cell cycle progression, proliferation, and tumor growth in prostate and liver cancers ^6, 7, 16^. To examine its role in spermatogonial cells, we first assessed the effects of CNNM1 knockdown in C18-4 cells; the cell cycle profile was altered, and the cells accumulated at G1/S phase. However, CNNM1 knockdown spermatogonial cells showed no significant difference in expression of apoptotic markers between CNNM1 knockdown cells and the control (unpublished data). Conversely, CNNM1 overexpression in C18-4 cells leads to an increase in the S-phase and G2/M phase population, reduction in G1and G0 population, indicative of enhanced proliferative activity. This proliferative shift was further supported by the upregulation of key proliferation markers, TOP2A and c-MYC, suggesting that CNNM1 promotes spermatogonial cell cycle progression and proliferation.

CNNM1 overexpression further enhanced the expression of c-*Myc, Klf4*, and TOP2A, key transcriptional regulators of proliferation, self-renewal, and telomerase activation. Both c-MYC and KLF4 activate TERT transcription, increasing telomerase activity—an essential feature of SSCs and undifferentiated spermatogonia ^17^. This finding aligns with reports of CNNM1-mediated promotion of cell proliferation in prostate and hepatocellular carcinomas ^6, 7^. Moreover, treatment with Ginsenoside Rh2 (GRh2), a CNNM1 inhibitor ^6^, reduced *Cnnm1* expression and induced apoptosis in spermatogonial cells, whereas CNNM1 overexpression mitigated this effect. Together, these data establish CNNM1 as a key determinant of spermatogonial proliferation and survival.

The global proteomic landscape of spermatogonial cells overexpressing CNNM1 was profiled using label-free quantitative proteomics. Gene ontology enrichment analysis revealed marked alterations in proteins associated with cell cycle regulation, metabolism, protein synthesis, and cellular stress responses—key pathways fundamental to SSC maintenance and self-renewal.

Among the upregulated proteins were GSTM1, YBX1, and several DEAD-box helicases (DDX1, DDX3X, DDX4, DDX5, DDX17, and DDX18). GSTM1, highly expressed in GFRA1-positive undifferentiated spermatogonia, contributes to SSC survival by modulating oxidative stress ^18^. YBX1, a multifunctional RNA/DNA-binding protein, is a known regulator of SSC maintenance and mediates GDNF-dependent translational control of transcripts linked to stemness and renewal ^19^. The upregulation of these proteins under CNNM1 overexpression supports a role for CNNM1 in reinforcing SSC identity and survival.

The concurrent induction of DEAD-box helicases further implicates CNNM1 in RNA metabolism and cell fate regulation. DDX1 and DDX5 are indispensable for SSC maintenance and proliferation ^20, 21^, while DDX3 regulates pluripotency and Wnt/β-catenin signaling ^8^. DDX4 is not expressed in PLZF-positive SSCs of the adult mouse testis, while it is highly expressed in in vitro cultured SSCs ^22^indicating a potential role under proliferative conditions. The coordinated upregulation of these helicases suggests that CNNM1 orchestrates transcriptional and post-transcriptional mechanisms that sustain germline stem cell renewal.

Additional proteins elevated by CNNM1 overexpression—Annexin A2 (ANXA2), Eif2s2, Prohibitin 2 (PHB2), and Keratin 8 (KRT8)—are likewise associated with proliferation and stem cell homeostasis. ANXA2 regulates cell division and maintains the integrity of the blood– testis barrier ^23^; EIF2S2 promotes proliferation through MYC stabilization and Wnt/β-catenin activation ^24^; PHB2 supports embryonic stem cell self-renewal ^25^; and KRT8 is a cytoskeletal marker linked to germ cell migration and morphogenesis ^26^. Their upregulation collectively underscores CNNM1’s role in sustaining an undifferentiated, proliferative SSC state.

Conversely, CNNM1 overexpression led to downregulation of AVPI, arginine vasopressin (VP), and Importin α5 (KPNA1)—proteins implicated in cell cycle arrest and differentiation ^27-29^. The suppression of these inhibitory regulators suggests that CNNM1 actively restrains differentiation cues to preserve SSC stemness.

Proteomic enrichment also revealed increased abundance of oxidoreductases, indicating that CNNM1 contributes to redox homeostasis. Because reactive oxygen species (ROS) can induce DNA damage and compromise SSC integrity ^30^ The activation of antioxidant pathways likely underlies CNNM1-mediated protection against oxidative stress. This mechanism is consistent with the need for robust redox regulation in germline stem cells undergoing continuous proliferation.

Collectively, these findings identify CNNM1 as a positive regulator of SSC proliferation and maintenance. CNNM1 expression is responsive to GDNF signaling and influences key cellular processes, including cell cycle progression, metabolic activity, and stress response. While the proteomic alterations observed here provide insight into CNNM1-associated molecular pathways, further mechanistic studies are required to delineate its direct targets and interaction partners. Overall, this work establishes CNNM1 as a component of the molecular framework that sustains SSC homeostasis and provides a basis for future investigations into its role in germline stem cell regulation and male fertility.

## Materials and methods

### Experimental animals

All animal procedures were approved by the Institutional Animal Ethics Committee of the Rajiv Gandhi Centre for Biotechnology, Thiruvananthapuram, India (IAEC/507/PRK/2015 and IAEC/776/PRK/2019). Mus musculus (Swiss albino strain) were maintained under specific pathogen-free conditions (22 ± 2 °C, 12 h light/dark cycle, 55 ± 5% humidity) with food and water ad libitum. Timed matings were set by overnight cohabitation of males and females, and the day of vaginal plug detection was designated embryonic day (E) 0.5. Plug-positive females were housed separately. Embryos were collected between E9.5 and E16.5; trunks (E9.5–E10.5) and genital ridges (E11.5–E16.5) were dissected for primordial germ cell analysis. Sex was determined morphologically from E12.5 onwards.

### cDNA synthesis and PCR

Total RNA was isolated using TRI Reagent (Sigma-Aldrich) and reverse-transcribed from 1 µg RNA with the High-Capacity cDNA Reverse Transcription Kit (Applied Biosystems). Conventional PCR was performed using gene-specific primers and Taq DNA polymerase (Thermo Fisher Scientific) under standard cycling conditions. Quantitative PCR was carried out using PowerUp SYBR Green Master Mix (Applied Biosystems) on a QuantStudio 6 Flex system, with Gapdh as the internal control. Relative expression was calculated by the ΔΔCt method. Details of the primer sets used are provided in Table S2.

### Plasmid Construction

The *Cnnm1* coding sequence (1.8 kb) was amplified from postnatal day 8 testicular cDNA using the primers Forward: 5′-CCGCTCGAGCGGATGGCGGCTGCCTTCCCG-3′ and Reverse: 5′-GGAATTCCTGTGATCAGGGGCGTTAAATTGGAG-3′. The PCR product and pEGFP-N1 vector were digested with XhoI and EcoRI, gel-purified, and ligated using T4 DNA ligase. The ligation mix was transformed into E. coli DH5α, and colonies were selected on kanamycin plates. Positive clones were verified by sequencing, and plasmids were purified using NucleoSpin kits (MACHEREY-NAGEL).

### Culture of C18-4 spermatogonial cell line

The mouse spermatogonial cell line C18-4 was maintained in Dulbecco’s MEM F12 (DMEM/F12) medium (Gibco) supplemented with 10% Fetal Bovine Serum (FBS) and 1% Antibiotic-Antimycotic at 37 °C in 5% CO_2_. Cells were seeded at 1.5–4 × 10^5^ cells cm^−2^ and sub cultured every 2–3 days upon reaching 70–75% confluence. For transfection, cells at ∼65% confluence were cultured in Opti-MEM and transfected with 2 µg plasmid DNA using Lipofectamine 2000 (Invitrogen) following the manufacturer’s instructions. GFP fluorescence was assessed 24 h post-transfection.

### Cell cycle analysis

C18-4 cells were harvested 24 h post-transfection, stained with 10 µg ml^−1^ Hoechst 33342 (Sigma-Aldrich) for 15 min at 37 °C, and analysed on a BD FACSAria flow cytometer (Becton Dickinson). GFP-positive populations were gated to determine cell-cycle distribution using FACS Diva software.

### GDNF treatment

Cells were cultured overnight in complete DMEM/F12 medium, then incubated for 24 h in serum-free medium containing 100 ng ml^−1^ glial cell line-derived neurotrophic factor (GDNF; Sigma-Aldrich). PBS served as the vehicle control. After treatment, cells were processed for immunofluorescence or RNA isolation. Experiments were performed in triplicate.

### Immunocytochemistry

C18-4 cells were grown on gelatin-coated coverslips, fixed with 4% paraformaldehyde for 15 min, permeabilized with 1% Triton X-100 for 10 min, and blocked with 5% BSA for 30 min at room temperature. Cells were incubated with primary antibodies overnight at 4 °C, followed by Alexa Fluor-conjugated secondary antibodies (Invitrogen) for 1 h in the dark. Nuclei were counterstained with DAPI (0.5 µg ml^−1^). Images were acquired using an Olympus FV3000 confocal laser-scanning microscope. Details of the antibodies used are provided in Table S3.

### Western blot analysis

Cells were lysed in RIPA buffer (25 mM Tris–HCl pH 7.6, 150 mM NaCl, 1% NP-40, 1% sodium deoxycholate, 0.1% SDS, and protease inhibitors; Roche). Lysates were cleared by centrifugation and quantified by the BCA assay. Proteins were separated by SDS–PAGE, transferred to PVDF membranes, blocked with 5% milk in PBST, and probed with primary and HRP-conjugated secondary antibodies. Proteins were visualized using diaminobenzidine/H_2_O_2_ and quantified by densitometry (GelDoc & Image Lab, Bio-Rad). Details of the antibodies used are provided in Table S3

### Proteomic analysis

Protein extracts were prepared in 50 mM ammonium bicarbonate containing RapiGest SF (Waters). Samples (100 µg) were reduced with dithiothreitol, alkylated with iodoacetamide, and digested overnight with trypsin (Sigma-Aldrich) at 37 °C. Peptides were analyzed on a nanoACQUITY UPLC coupled to a SYNAPT-G2 Q-TOF mass spectrometer (Waters). Peptide separation was achieved on a BEH C18 column (75 µm × 100 mm, 1.7 µm) using a linear 1– 40% acetonitrile gradient over 55 min at 300 nl min^−1^. Data were acquired in MSE mode with lock-mass correction using [Glu^1^]-fibrinopeptide B.

Spectra were processed with Progenesis QI for Proteomics (Nonlinear Dynamics) using the Mus musculus UniProt database. Carbamidomethylation of cysteine and oxidation of methionine were set as fixed and variable modifications, respectively. FDR was controlled at <4%. Label-free quantification was performed using normalized intensities from three biological replicates. Proteins with fold-change > 1.5 or < 0.5 (p < 0.05, Student’s t-test) were considered significantly altered. Functional enrichment was assessed using GONet and PANTHER.

### Statistics

Statistical analyses were performed using GraphPad Prism, and P value of 0.05 or below was considered significant.

## Acknowledgements

This work was supported by Grants SR/SO/HS-140/2012 from Department of Science and Technology, Government of India, New Delhi; CRG/2020/000649 from Science and Engineering Research Board, Government of India, New Delhi; DBT/HRD/34/07/2007 from Department of Biotechnology, Government of India, New Delhi; 5/10/FR/47/2020-RBMCH from Indian Council of Medical Research, New Delhi; Intramural funding from Rajiv Gandhi Centre for Biotechnology, Thiruvananthapuram and Emeritus Scientist Award No. HRD/Head/IES/2023 from Indian Council of Medical Research, New Delhi to PGK. IK was supported by Grant No. DST/INSPIRE Fellowship/2015/IF150354 by Department of Science and Technology, Government of India, New Delhi. Mahesh Kumar and Arun Surendran at the Mass Spectrometry and Proteomics Core Facility assisted in 2D LC-MS/MS analysis.

## Author Contributions

PKG conceptualised the study, secured funding, designed the experiment and supervised the study. and edited the manuscript. IK performed the experiments, analysed and interpreted the data, and wrote the manuscript.

## Figure Legends

**Figure S1| CNNM1 overexpression in C18-4 cells**. Immunofluorescence analysis of C18-4 cells transfected with pEGFPN1 (control) or Cnnm1-pEGFPN1. GFP (green) indicates transfected cells, CNNM1 (red) shows CNNM1 localization, and DAPI (blue) stains nuclei. Merged images (yellow) indicate colocalization of CNNM1 with GFP. Scale bar = 20 μm (A); Schematic representation of the Cnnm1 overexpression construct (B); Western blot analysis showing overexpression of the CNNM1-GFP fusion protein (C); Influence of CNNM1 overexpression on gene expression analyzed by RT-PCR. Expression levels of Top2a and Itgb1 transcripts were evaluated in three biological replicates (D).

**Figure S2| Canonical pathways associated with upregulated proteins in CNNM1-overexpressed C18-4 cells**. Pathway analysis was performed using PANTHER, and proteins with fold change > 1.5 were considered upregulated. The figure shows canonical pathways enriched among these upregulated proteins.

