## Supplementary figures and images for "CNNM1 is Involved in Spermatogonial Stem Cell Maintenance in the Mouse"

### Supplemental Figures S1-S2

**Figure S1**

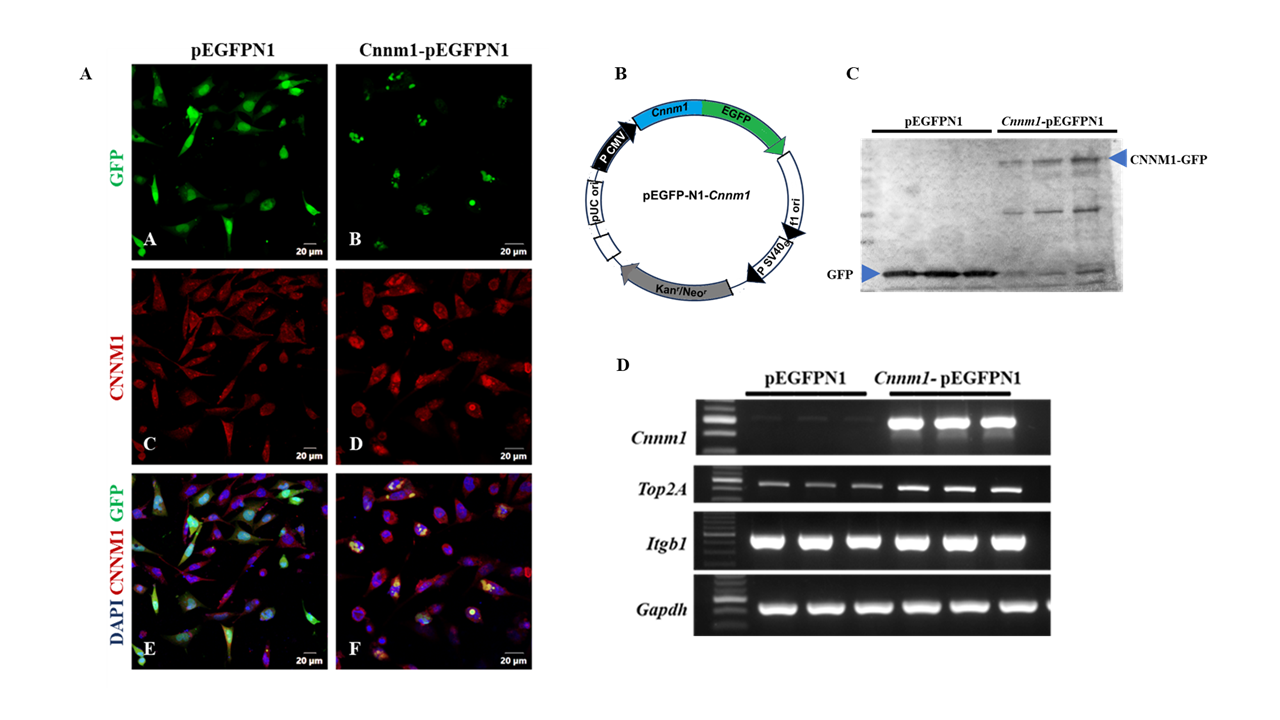

**Figure S2**

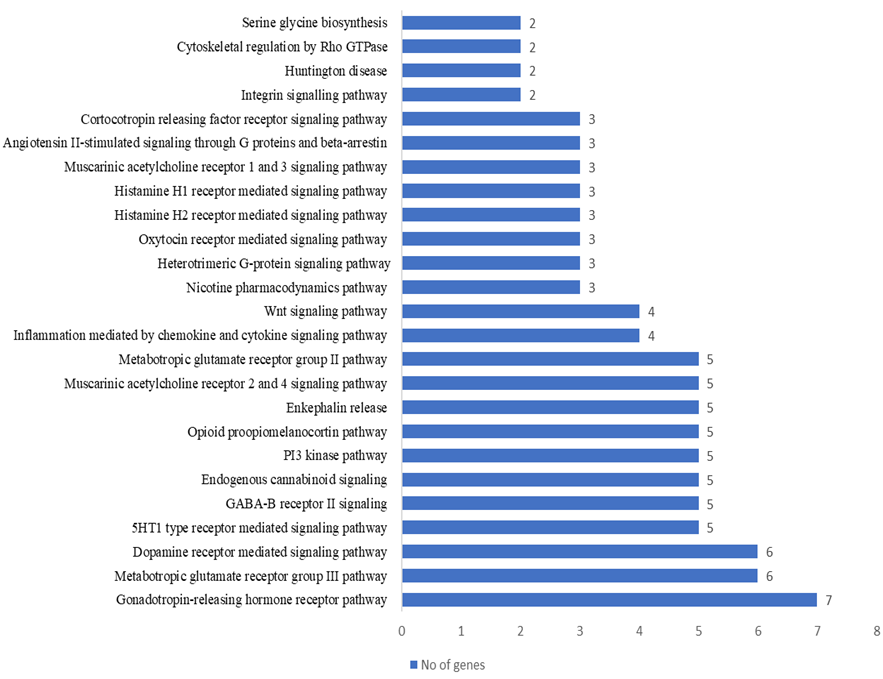
