## Supplemental Table S1 for "CNNM1 is Involved in Spermatogonial Stem Cell Maintenance in the Mouse"

| **Uniprot ID** | **Gene Symbol** | **Protein description** | **Fold change** |
| --- | --- | --- | --- |
| **Q6ZQ38** | *Cand1* | Cullin-associated NEDD8-dissociated protein 1 | 2.88 |
| **P57780** | *Actn4* | Alpha-actinin-4 | 1.53 |
| **Q61656** | *Ddx5* | Probable ATP-dependent RNA helicase DDX5 | 1.84 |
| **Q8VDJ3** | *Hdlbp* | Vigilin | 1.99 |
| **O70133** | *Dhx9* | ATP-dependent RNA helicase A | 1.60 |
| **P42932** | *Cct8* | T-complex protein 1 subunit theta | 1.83 |
| **P68372** | *Tubb4b* | Tubulin beta-4B chain | 1.56 |
| **P99024** | *Tubb5* | Tubulin beta-5 chain | 1.51 |
| **Q920B9** | *Supt16h* | FACT complex subunit SPT16 | 2.96 |
| **P62908** | *Rps3* | 40S ribosomal protein S3 | 2.36 |
| **P62702** | *Rps4x* | 40S ribosomal protein S4 | 1.90 |
| **Q01320** | *Top2a* | DNA topoisomerase 2-alpha | 2.33 |
| **Q9D0E1** | *Hnrnpm* | Heterogeneous nuclear ribonucleoprotein M | 2.24 |
| **Q91YQ5** | *Rpn1* | Dolichyl-diphosphooligosaccharide--protein glycosyltransferase subunit 1 | 2.65 |
| **P51881** | *Slc25a5* | ADP/ATP translocase 2 | 1.88 |
| **Q9D6Z1** | *Nop56* | Nucleolar protein 56 | 1.88 |
| **P08249** | *Mdh2* | Malate dehydrogenase | 2.84 |
| **Q6ZWN5** | *Rps9* | 40S ribosomal protein S9 | 2.13 |
| **P97351** | *Rps3a* | 40S ribosomal protein S3a | 1.93 |
| **Q8BMK4** | *Ckap4* | Cytoskeleton-associated protein 4 | 2.03 |
| **O88569** | *Hnrnpa2b1* | Heterogeneous nuclear ribonucleoproteins A2/B1 | 1.73 |
| **Q921M3** | *Sf3b3* | Splicing factor 3B subunit 3 | 1.86 |
| **P07356** | *Anxa2* | Annexin A2 | 1.73 |
| **Q501J6** | *Ddx17* | Probable ATP-dependent RNA helicase DDX17 | 2.12 |
| **P27659** | *Rpl3* | 60S ribosomal protein L3 | 2.49 |
| **P25444** | *Rps2* | 40S ribosomal protein S2 | 2.10 |
| **Q62167** | *Ddx3x* | ATP-dependent RNA helicase DDX3X | 1.95 |
| **P12970** | *Rpl7a* | 60S ribosomal protein L7a | 1.71 |
| **A2AQ07** | *Tubb1* | Tubulin beta-1 chain | 1.85 |
| **Q9ESZ8** | *Gtf2i* | General transcription factor II-I | 1.55 |
| **Q61937** | *Npm1* | Nucleophosmin | 1.62 |
| **P62806** | *H4c1* | Histone H4 | 6.24 |
| **P02301** | *H3-5* | Histone H3.3C | 5.34 |
| **Q62318** | *Trim28* | Transcription intermediary factor 1-beta | 2.11 |
| **P49312** | *Hnrnpa1* | Heterogeneous nuclear ribonucleoprotein A1 | 1.53 |
| **Q99JY0** | *Hadhb* | Trifunctional enzyme subunit beta | 1.62 |
| **P50580** | *Pa2g4* | Proliferation-associated protein 2G4 | 1.52 |
| **P48962** | *Slc25a4* | ADP/ATP translocase 1 | 1.54 |
| **Q9D3R6** | *Katnal2* | Katanin p60 ATPase-containing subunit A-like 2 | 1.55 |
| **P11679** | *Krt8* | Keratin | 2.05 |
| **P07742** | *Rrm1* | Ribonucleoside-diphosphate reductase large subunit | 2.44 |
| **O88342** | *Wdr1* | WD repeat-containing protein 1 | 1.77 |
| **Q9CZN7** | *Shmt2* | Serine hydroxymethyltransferase | 1.57 |
| **P10854** | *H2bc14* | Histone H2B type 1-M | 3.74 |
| **Q6R0H7** | *Gnas* | Guanine nucleotide-binding protein G | 2.65 |
| **A6PWD2** | *Fhad1* | Forkhead-associated domain-containing protein 1 | 2.54 |
| **Q9Z0N1** | *Eif2s3x* | Eukaryotic translation initiation factor 2 subunit 3 | 2.40 |
| **A3KGS3** | *Ralgapa2* | Ral GTPase-activating protein subunit alpha-2 | 1.79 |
| **Q91VR5** | *Ddx1* | ATP-dependent RNA helicase DDX1 | 1.90 |
| **P53026** | *Rpl10a* | 60S ribosomal protein L10a | 1.83 |
| **P97461** | *Rps5* | 40S ribosomal protein S5 [Cleaved into: 40S ribosomal protein S5 | 2.06 |
| **P63276** | *Rps17* | 40S ribosomal protein S17 | 1.73 |
| **Q8BG05** | *Hnrnpa3* | Heterogeneous nuclear ribonucleoprotein A3 | 1.99 |
| **P62281** | *Rps11* | 40S ribosomal protein S11 | 1.98 |
| **P19253** | *Rpl13a* | 60S ribosomal protein L13a | 1.58 |
| **Q7TMK9** | *Syncrip* | Heterogeneous nuclear ribonucleoprotein Q | 1.92 |
| **P62301** | *Rps13* | 40S ribosomal protein S13 | 2.18 |
| **Q9CZX8** | *Rps19* | 40S ribosomal protein S19 | 1.95 |
| **P10649** | *Gstm1* | Glutathione S-transferase Mu 1 | 1.65 |
| **Q9DBH5** | *Lman2* | Vesicular integral-membrane protein VIP36 | 1.75 |
| **Q9CZM2** | *Rpl15* | 60S ribosomal protein L15 | 1.53 |
| **Q3UV17** | *Krt76* | Keratin | 2.98 |
| **Q9Z1Q5** | *Clic1* | Chloride intracellular channel protein 1 | 1.72 |
| **P51150** | *Rab7a* | Ras-related protein Rab-7a | 2.13 |
| **P14131** | *Rps16* | 40S ribosomal protein S16 | 1.95 |
| **P62082** | *Rps7* | 40S ribosomal protein S7 | 2.14 |
| **Q99KP6** | *Prpf19* | Pre-mRNA-processing factor 19 | 1.55 |
| **P50396** | *Gdi1* | Rab GDP dissociation inhibitor alpha | 1.85 |
| **Q8VEM8** | *Slc25a3* | Phosphate carrier protein | 1.78 |
| **Q922Q8** | *Lrrc59* | Leucine-rich repeat-containing protein 59 [Cleaved into: Leucine-rich repeat-containing protein 59 | 1.89 |
| **P67778** | *Phb1* | Prohibitin 1 | 1.86 |
| **Q8VDN2** | *Atp1a1* | Sodium/potassium-transporting ATPase subunit alpha-1 | 2.02 |
| **P62754** | *Rps6* | 40S ribosomal protein S6 | 1.72 |
| **P63037** | *Dnaja1* | DnaJ homolog subfamily A member 1 | 1.61 |
| **P47963** | *Rpl13* | 60S ribosomal protein L13 | 1.78 |
| **P68433** | *H3c1* | Histone H3.1 | 6.72 |
| **P84228** | *H3c13* | Histone H3.2 | 6.54 |
| **P62137** | *Ppp1ca* | Serine/threonine-protein phosphatase PP1-alpha catalytic subunit | 1.53 |
| **Q8R1M2** | *H2aj* | Histone H2A.J | 6.01 |
| **Q8VH51** | *Rbm39* | RNA-binding protein 39 | 1.90 |
| **P46664** | *Adss2* | Adenylosuccinate synthetase isozyme 2 | 1.79 |
| **Q78ZA7** | *Nap1l4* | Nucleosome assembly protein 1-like 4 | 1.55 |
| **O88544** | *Cops4* | COP9 signalosome complex subunit 4 | 1.92 |
| **Q8VIJ6** | *Sfpq* | Splicing factor | 1.72 |
| **P46638** | *Rab11b* | Ras-related protein Rab-11B | 1.75 |
| **Q9Z204** | *Hnrnpc* | Heterogeneous nuclear ribonucleoproteins C1/C2 | 1.63 |
| **P53994** | *Rab2a* | Ras-related protein Rab-2A | 1.52 |
| **P62264** | *Rps14* | 40S ribosomal protein S14 | 1.97 |
| **Q8BMD8** | *Slc25a24* | Calcium-binding mitochondrial carrier protein SCaMC-1 | 1.95 |
| **P61027** | *Rab10* | Ras-related protein Rab-10 | 2.25 |
| **P43277** | *H1-3* | Histone H1.3 | 1.99 |
| **Q61496** | *Ddx4* | ATP-dependent RNA helicase DDX4 | 1.96 |
| **O35129** | *Phb2* | Prohibitin-2 | 3.68 |
| **P07744** | *Krt4* | Keratin | 1.77 |
| **Q8K363** | *Ddx18* | ATP-dependent RNA helicase DDX18 | 2.06 |
| **Q9DB15** | *Mrpl12* | 39S ribosomal protein L12 | 1.61 |
| **Q8R050** | *Gspt1* | Eukaryotic peptide chain release factor GTP-binding subunit ERF3A | 1.97 |
| **Q9DBG6** | *Rpn2* | Dolichyl-diphosphooligosaccharide--protein glycosyltransferase subunit 2 | 1.75 |
| **P62849** | *Rps24* | 40S ribosomal protein S24 | 1.72 |
| **P50431** | *Shmt1* | Serine hydroxymethyltransferase | 3.16 |
| **Q9CQM9** | *Glrx3* | Glutaredoxin-3 | 1.81 |
| **P51410** | *Rpl9* | 60S ribosomal protein L9 | 2.64 |
| **P62717** | *Rpl18a* | 60S ribosomal protein L18a | 2.20 |
| **P84099** | *Rpl19* | 60S ribosomal protein L19 | 2.06 |
| **Q91ZW3** | *Smarca5* | SWI/SNF-related matrix-associated actin-dependent regulator of chromatin subfamily A member 5 | 1.78 |
| **P55821** | *Stmn2* | Stathmin-2 | 1.72 |
| **P23198** | *Cbx3* | Chromobox protein homolog 3 | 2.72 |
| **Q91VR2** | *Atp5f1c* | ATP synthase subunit gamma | 1.98 |
| **P62900** | *Rpl31* | 60S ribosomal protein L31 | 1.90 |
| **Q9DB05** | *Napa* | Alpha-soluble NSF attachment protein | 1.63 |
| **Q8CGK7** | *Gnal* | Guanine nucleotide-binding protein G | 1.52 |
| **O08810** | *Eftud2* | 116 kDa U5 small nuclear ribonucleoprotein component | 2.94 |
| **P61255** | *Rpl26* | 60S ribosomal protein L26 | 1.63 |
| **Q9CQE8** | *RTRAF* | RNA transcription | 1.63 |
| **Q9WV02** | *Rbmx* | RNA-binding motif protein | 2.08 |
| **P35980** | *Rpl18* | 60S ribosomal protein L18 | 1.99 |
| **Q9CQA3** | *Sdhb* | Succinate dehydrogenase [ubiquinone] iron-sulfur subunit | 2.17 |
| **Q9D1G1** | *Rab1b* | Ras-related protein Rab-1B | 1.60 |
| **A2AEY4** | *Map7d3* | MAP7 domain-containing protein 3 | 1.88 |
| **Q8BHC4** | *Dcakd* | Dephospho-CoA kinase domain-containing protein | 1.78 |
| **A2RSX7** | *Tyw5* | tRNA wybutosine-synthesizing protein 5 | 2.52 |
| **Q8R480** | *Nup85* | Nuclear pore complex protein Nup85 | 1.71 |
| **P61971** | *Nutf2* | Nuclear transport factor 2 | 1.67 |
| **P35979** | *Rpl12* | 60S ribosomal protein L12 | 1.65 |
| **P62960** | *Ybx1* | Y-box-binding protein 1 | 2.08 |
| **Q9CQQ7** | *Atp5pb* | ATP synthase F | 1.88 |
| **P14115** | *Rpl27a* | 60S ribosomal protein L27a | 1.66 |
| **P60867** | *Rps20* | 40S ribosomal protein S20 | 2.29 |
| **P99027** | *Rplp2* | 60S acidic ribosomal protein P2 | 1.94 |
| **Q9D1R9** | *Rpl34* | 60S ribosomal protein L34 | 1.64 |
| **P20108** | *Prdx3* | Thioredoxin-dependent peroxide reductase | 1.63 |
| **P62889** | *Rpl30* | 60S ribosomal protein L30 | 2.26 |
| **P43275** | *H1-1* | Histone H1.1 | 1.80 |
| **P56564** | *Slc1a3* | Excitatory amino acid transporter 1 | 2.82 |
| **P62852** | *Rps25* | 40S ribosomal protein S25 | 2.23 |
| **Q64096** | *Mcf2l* | Guanine nucleotide exchange factor DBS | 2.03 |
| **P15532** | *Nme1* | Nucleoside diphosphate kinase A | 1.88 |
| **P59764** | *Dock4* | Dedicator of cytokinesis protein 4 | 1.77 |
| **P18155** | *Mthfd2* | Bifunctional methylenetetrahydrofolate dehydrogenase/cyclohydrolase | 1.55 |
| **Q60932** | *Vdac1* | Voltage-dependent anion-selective channel protein 1 | 4.27 |
| **Q80X50** | *Ubap2l* | Ubiquitin-associated protein 2-like | 3.24 |
| **O54734** | *Ddost* | Dolichyl-diphosphooligosaccharide--protein glycosyltransferase 48 kDa subunit | 2.88 |
| **P0C0S6** | *H2az1* | Histone H2A.Z | 2.52 |
| **Q99L45** | *Eif2s2* | Eukaryotic translation initiation factor 2 subunit 2 | 1.98 |
| **P43276** | *H1-5* | Histone H1.5 | 1.64 |
| **Q8BL97** | *Srsf7* | Serine/arginine-rich splicing factor 7 | 1.95 |
| **P83882** | *Rpl36a* | 60S ribosomal protein L36a | 1.61 |
| **Q6PDM2** | *Srsf1* | Serine/arginine-rich splicing factor 1 | 1.56 |
| **Q9CX86** | *Hnrnpa0* | Heterogeneous nuclear ribonucleoprotein A0 | 1.51 |
| **P07091** | *S100a4* | Protein S100-A4 | 42.50 |
| **Q9D0M1** | *Prpsap1* | Phosphoribosyl pyrophosphate synthase-associated protein 1 | 3.11 |
| **Q9D6I7** | *Dipk1a* | Divergent protein kinase domain 1A | 2.36 |
| **Q569L8** | *Cenpj* | Centromere protein J | 1.95 |
| **Q64518** | *Atp2a3* | Sarcoplasmic/endoplasmic reticulum calcium ATPase 3 | 1.90 |
| **Q9CXG3** | *Ppil4* | Peptidyl-prolyl cis-trans isomerase-like 4 | 1.79 |
| **Q80Z25** | *Ofd1* | Oral-facial-digital syndrome 1 protein homolog | 1.69 |
| **Q9CQI9** | *Med30* | Mediator of RNA polymerase II transcription subunit 30 | 5.83 |
| **O70503** | *Hsd17b12* | Very-long-chain 3-oxoacyl-CoA reductase | 2.26 |
| **Q9DC51** | *Gnai3* | Guanine nucleotide-binding protein G | 2.24 |
| **P62880** | *Gnb2* | Guanine nucleotide-binding protein G | 2.15 |
| **P61082** | *Ube2m* | NEDD8-conjugating enzyme Ubc12 | 1.86 |
| **P25688** | *Uox* | Uricase | 1.80 |
| **Q9Z1W8** | *Atp12a* | Potassium-transporting ATPase alpha chain 2 | 1.77 |
| **P62855** | *Rps26* | 40S ribosomal protein S26 | 1.77 |
| **Q9CQW2** | *Arl8b* | ADP-ribosylation factor-like protein 8B | 1.61 |
| **Q9QXL1** | *Kif21b* | Kinesin-like protein KIF21B | 1.54 |
| **Q8C650** | *Septin10* | Septin-10 | 2.49 |
| **Q6ZWV7** | *Rpl35* | 60S ribosomal protein L35 | 2.44 |
| **P61514** | *Rpl37a* | 60S ribosomal protein L37a | 1.97 |
| **Q9CR57** | *Rpl14* | 60S ribosomal protein L14 | 1.95 |
| **Q8VHE0** | *Sec63* | Translocation protein SEC63 homolog | 1.50 |
| **B2RSH2** | *Gnai1* | Guanine nucleotide-binding protein G | 2.62 |
| **Q8K0E1** | *Kctd15* | BTB/POZ domain-containing protein KCTD15 | 2.41 |
| **Q9DCN2** | *Cyb5r3* | NADH-cytochrome b5 reductase 3 | 2.39 |
| **Q9JKK1** | *Stx6* | Syntaxin-6 | 2.18 |
| **P34884** | *Mif* | Macrophage migration inhibitory factor | 2.16 |
| **Q9DAW9** | *Cnn3* | Calponin-3 | 2.05 |
| **O35435** | *Dhodh* | Dihydroorotate dehydrogenase | 1.72 |
| **Q8R1F5** | *Hyi* | Putative hydroxypyruvate isomerase | 1.69 |
| **Q8BRM2** | *Gorab* | RAB6-interacting golgin | 1.55 |
| **Q60931** | *Vdac3* | Voltage-dependent anion-selective channel protein 3 | 1.53 |
| **P24527** | *Lta4h* | Leukotriene A-4 hydrolase | 1.76 |
| **P35293** | *Rab18* | Ras-related protein Rab-18 | 1.52 |
| **P84089** | *Erh* | Enhancer of rudimentary homolog | 1.52 |
| **P09671** | *Sod2* | Superoxide dismutase [Mn] | 2.48 |
| **P59999** | *Arpc4* | Actin-related protein 2/3 complex subunit 4 | 1.74 |
| **P61759** | *Vbp1* | Prefoldin subunit 3 | 1.62 |
| **Q9D0T1** | *Snu13* | NHP2-like protein 1 | 1.56 |
| **Q7TQ95** | *Lnpk* | Endoplasmic reticulum junction formation protein lunapark | 3.58 |
| **Q64437** | *Adh7* | All-trans-retinol dehydrogenase [NAD | 3.51 |
| **P35283** | *Rab12* | Ras-related protein Rab-12 | 3.11 |
| **Q01730** | *Rsu1* | Ras suppressor protein 1 | 2.89 |
| **P09023** | *Hoxb6* | Homeobox protein Hox-B6 | 2.24 |
| **Q7SIG6** | *Asap2* | Arf-GAP with SH3 domain | 2.18 |
| **Q9D312** | *Krt20* | Keratin | 2.17 |
| **Q9D880** | *Timm50* | Mitochondrial import inner membrane translocase subunit TIM50 | 2.16 |
| **O35326** | *Srsf5* | Serine/arginine-rich splicing factor 5 | 1.73 |
| **Q9CX47** | *Xrcc2* | DNA repair protein XRCC2 | 1.71 |
| **Q64261** | *Cdk6* | Cyclin-dependent kinase 6 | 1.68 |
| **P60766** | *Cdc42* | Cell division control protein 42 homolog | 1.65 |
| **Q91V41** | *Rab14* | Ras-related protein Rab-14 | 1.50 |
| **Q9CQX2** | *Cyb5b* | Cytochrome b5 type B | 3.30 |
| **Q9CR67** | *Tmem33* | Transmembrane protein 33 | 3.26 |
| **Q9CPR8** | *Nsmce3* | Non-structural maintenance of chromosomes element 3 homolog | 2.73 |
| **Q60930** | *Vdac2* | Voltage-dependent anion-selective channel protein 2 | 2.38 |
| **Q9DCX2** | *Atp5pd* | ATP synthase subunit d | 2.05 |
| **P59325** | *Eif5* | Eukaryotic translation initiation factor 5 | 1.63 |
| **Q6P5F9** | *Xpo1* | Exportin-1 | 1.60 |
| **P28658** | *Atxn10* | Ataxin-10 | 1.57 |
| **P50543** | *S100a11* | Protein S100-A11 | 1.55 |
| **O35618** | *Mdm4* | Protein Mdm4 | 1.50 |
| **P11983** | *Tcp1* | T-complex protein 1 subunit alpha | 1.89 |
| **P80315** | *Cct4* | T-complex protein 1 subunit delta | 1.52 |
| **P80314** | *Cct2* | T-complex protein 1 subunit beta | 1.65 |
| **P80318** | *Cct3* | T-complex protein 1 subunit gamma | 1.54 |
| **Q8CG76** | *Akr7a2* | Aflatoxin B1 aldehyde reductase member 2 | 0.24 |
| **Q9D7H4** | *Avpi1* | Arginine vasopressin-induced protein 1 | 0.26 |
| **Q60960** | *Kpna1* | Importin subunit alpha-5 | 0.26 |
| **P51855** | *Gss* | Glutathione synthetase | 0.30 |
| **Q9QXB9** | *Drg2* | Developmentally-regulated GTP-binding protein 2 | 0.37 |
| **Q9JHI5** | *Ivd* | Isovaleryl-CoA dehydrogenase | 0.37 |
| **Q9CPT4** | *Mydgf* | Myeloid-derived growth factor | 0.40 |
| **Q8BGB8** | *Coq4* | Ubiquinone biosynthesis protein COQ4 homolog | 0.45 |
| **Q8CIC2** | *Nup42* | Nucleoporin NUP42 | 0.46 |
| **Q9D110** | *Mthfs* | 5-formyltetrahydrofolate cyclo-ligase | 0.46 |
| **Q6IFZ9** | *Krt74* | Keratin | 0.48 |
| **Q64737** | *Gart* | Trifunctional purine biosynthetic protein adenosine-3 [Includes: Phosphoribosylamine--glycine ligase | 0.50 |
| **Q91V12** | *Acot7* | Cytosolic acyl coenzyme A thioester hydrolase | 0.50 |

**Table S1 List of differentially displayed proteins from proteome profiling of CNNM1 overexpression in C18-4 spermatogonial cells.** Proteins with fold change ≥ 1.5 are considered to be upregulated and ≤ 0.5 as downregulated.
