## Supplemental Table S2-S3 for "CNNM1 is Involved in Spermatogonial Stem Cell Maintenance in the Mouse"

**Sequences of primers used in the RT-PCR experiment**

| **Sl. No** | **Gene** | **Sequence (5′ to 3′)** | **Type** |
| --- | --- | --- | --- |
| 1. | *Gapdh* | TCACCACCATGGAGAAGGC | Forward |
|  |  | GCTAAGCAGTTGGTGGTGCA | Reverse |
| 2. | *Top2a* | CAACTGGAACATATACTGCTCCG | Forward |
|  |  | GGGTCCCTTTGTTTGTTATCAGC | Reverse |
| 3. | *Itgb1* | TGTGGACAGTGTGTGTGTAGGA | Forward |
|  |  | GGACCAGTAGGACAGTCTGGAG | Reverse |
| 4. | *Cnnm1* | AGGTGATGGGCATTGTGACT | Forward |
|  |  | CCTTCCGCACGTCATTATC | Reverse |
| 5. | Cnnm1-cloning primers | CCGCTCGAGCGGATGGCGGCTGCCTTCCCG | Forward |
|  |  | GGAATTCCTGTGATCAGGGGCGTTAAATTGGAG | Reverse |

**Table S3**

Antibodies used in the study

| **Sl. No.** | **Antibody** | **Vendor** | **Type** |
| --- | --- | --- | --- |
| 1. | GAPDH | Santa cruz | **Primary** |
| 2. | CNNM1 | In House |  |
| 3. | TOP2A | Santa Cruz |  |
| 4. | MIF | Abcam |  |
| 5. | GFP | Santa Cruz |  |
| 6. | DDX3 | Santa Cruz |  |
| 7. | DNAJA1 | Abcam |  |
| 8. | DBPA | Abcam |  |
| 9. | Donkey anti mouse IgG conjugated with horse radish peroxidase | Santa Cruz | **Secondary** |
| 10. | Goat anti rabbit IgG conjugated with horse radish peroxidase | Santa Cruz |  |
| 11. | Rabbit anti goat IgG conjugated with horse radish peroxidase | Santa Cruz |  |
| 12. | Alexa fluor 488 rabbit IgG (H+L) antibody | Invitrogen |  |
| 13. | Alexa fluor 568 rabbit IgG (H+L) antibody | Invitrogen |  |
